# Is Melanopsin Activation Affecting Large Field Color Matching Functions?

**DOI:** 10.1101/2022.03.11.484009

**Authors:** Pablo Barrionuevo, Clemente Paz Filgueira, Dingcai Cao

## Abstract

Color theory is based on the exclusive activation of cones. However, since the discovery of melanopsin expressing cells in the human retina, evidence of its intrusion in brightness and color vision is increasing. We aimed to assess if differences between peripheral or large field and foveal color matches can be accounted for melanopsin activation or rod intrusion. Photopic color matches by young observers showed that differences between extrafoveal and foveal results cannot be explained by rod intrusion. Furthermore, statistical analyses on existing color matching functions suggest a role of melanopsin activation, particularly, in Large Field *S* Fundamentals.

## Introduction

At photopic light levels, any light can be color-matched by mixing three different colored lights (i.e. primaries), due to trichromacy of human color vision mediated by three types of cones (L-, M- and S- cones). Color matching functions (CMFs) represent the relative contributions of the primaries to match monochromatic test lights in the visible spectrum. Based on CMFs, color specification for any visible light can be achieved by metrics called colorimetry. For example, the CIE colorimetric system, developed since 1931 (CIE, 1932, 1964), comprises “the essential standards and procedures of measurement that are necessary to make colorimetry a useful tool in science and technology” (Wyszecki & Stiles, 2000). This system is based on human vision and it proposes an objective notation, independent of idiosyncratic use of color names (Nakano, 1996). However, photopic color matches depend on the field of view. For larger stimuli incorporating the perifoveal visual field, normal color perception was not satisfied by the 2° colorimetric standard observer. Peripheral color matching data produced a desaturated appearance compared with foveal perception (Moreland & Cruz, 1959). Therefore, a new set of CMFs for large field 10° field size was proposed by CIE (1964), for applications having stimuli larger than 4° of visual field. The differences between the small and large field CMFs were attributed to the macular pigment (CIE, 2006) or rod intrusion (Palmer, 1981; Trezona, 1970).

*L*, *M* and *S* cone fundamentals (Smith & Pokorny, 1975), were obtained by linear transformations of traditional CMFs, for example, 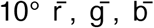 from Stiles and Burch (1959). However, since the discovery of melanopsin expressing intrinsic photosensitive retinal ganglion cells (ipRGCs) in the peripheral retina (Berson et al., 2002; Dacey et al., 2005; Hattar et al., 2002), several works have suggested the involvement of melanopsin activation in color vision. IpRGCs projections include centers for non-image forming functions, but also the Lateral Geniculate Nucleus (LGN) and superior colliculus that are involved in image-forming visual processes (Hattar et al., 2003). Melanopsin is a photopigment with a peak around 480 nm (Provencio et al., 2000). Its contribution to non-image forming processes such as pupillary light reflex and circadian photoentrainment is well characterized (Aranda & Schmidt, 2020; Foster et al., 2020; Spitschan, 2019), while evidence of a role in image-forming functions is being accumulated in the last years (Lucas et al., 2020). It was shown that human brightness perception is affected by melanopsin activation (Brown et al., 2012; Yamakawa et al., 2019; Zele, Adhikari, et al., 2018), and it was suggested that melanopsin could be involved in color constancy (Barrionuevo & Cao, 2019), unique white perception (Cao et al., 2018), color threshold (Horiguchi et al., 2013), and color processing (Zele, Feigl, et al., 2018). In this study, we aimed to assess if differences between peripheral or large field and foveal color matches can be accounted by melanopsin activation and/or rod intrusion.

## Methods

### Participants

Four participants (3 males and 1 female) participated in this study (37 ± 8.2 years old). All participants had normal color vision assessed by the Farnsworth-Munsell 100 Hue test and the Nagel anomaloscope. Three of the participants were experienced in psychophysical experiments and one participant was naïve subject. The study protocols were approved by the Institutional Review Board at the University of Illinois at Chicago and were in compliance with the Declaration of Helsinki.

### Apparatus

We developed a color matching optical system based on two optical branches, one for the matching light and the other for the test light (Fig. 1A). The matching light branch consisted of a set of three primaries [Fig. 1B, peak wavelengths: 460 nm (B), 520 nm (G) and 630 nm (R)] formed by intensity-adjustable bright LEDs and interference filters. The lights from the primaries were combined through a custom-made fiber optics bundle and a homogenizer (Cao et al., 2015). The test light branch consisted of a xenon lamp, UV-blocking filters, an electronic shutter, and a filter holder to locate spectral and neutral density filters. Spectral filters covered the range from 470 nm to 620 nm with full width at half maximum of 10 nm (Figs. 1C and 1D). Test lights with shorter peak wavelengths than 470 nm were not tested because the light levels were too low. Further, test lights with wavelengths longer than 620 nm were excluded as melanopsin activation in these wavelengths is negligible. Neutral density filters were used to help to maintain a relatively constant light level (mean =1780 td ± s. d.= 520 td) for all test lights. Both branches formed circular light patches that were overlapped using a semitransparent glass in order to stimulate the same retinal area. A field lens with a 2 mm artificial pupil was used to create a Maxwellian view.

**Fig. 1.**
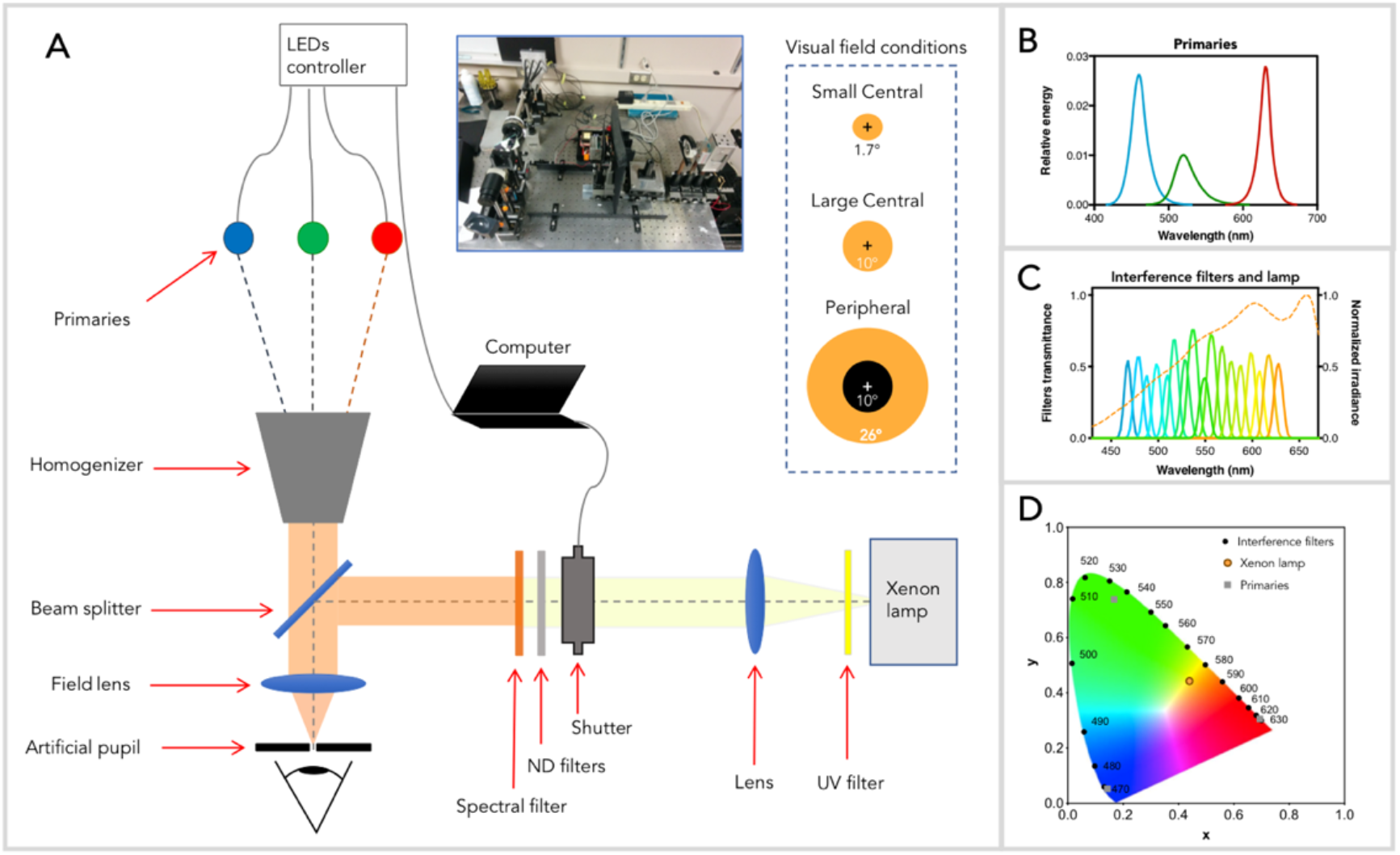
Experimental setup. A) Schematic representation of the optical arrangements used to obtain the color matches. A photograph of the actual arrangement is shown in the inset (it was covered with a black box during experiment sessions). Spatial dispositions of the visual field conditions are represented as insets on the right of the panel. Spectral and chromaticity values of matching (B and D) and test (C and D) fields are shown.

Presentation of both stimuli was sequential, three seconds each. The temporal sequence was obtained producing a square wave for the matching stimulus in counter phase to the trigger for opening the mechanical shutter of the test stimulus.

During the matching period the participant could vary the intensity of each primary using a gamepad. For short-wavelength test lights, the R primary was combined with the test stimulus instead of the matching stimulus. The stimulus presentation was controlled by self-developed software on Objective-C on a 27” iMac Apple computer.

### Procedure

The measurements were conducted in three spatial visual field conditions: 1) “Peripheral”, consisting in an annular field with inner edge diameter of 10° and outer edge diameter of 26°; 2) “Large Central”, consisting in a circular field of 10° diameter; and 3) “Small Central”, consisting in a circular field of 1.7° diameter.

During each trial, each participant fixated at the center of the stimulus area and adjusted the intensity of each primary of the matching stimulus in order to match the color of the test stimulus. Sequential presentation was maintained until the participant considered that a satisfactory match was found. Each observer obtained at least three matches for each test stimulus per condition.

The Peripheral condition was repeated after bleaching in order to minimize rods contribution. Participants were exposed during 15 s. to ~160000 sc. td provided by the Xenon lamp after UV blocking and neutral density filtering of 1 log unit. This light exposure bleached the 85% of rod photopigment approximately (Geisler & Banks, 2010). After bleaching, the participant waited 120 s in darkness to recover cone sensitivity. This bleaching sequence was repeated before each test wavelength. The bleaching experiment was carried out for test stimuli from 470 nm – 530 nm.

The outputs of each match were the relatively intensity values for *R*, *G* and *B* primaries. The *R*, *G* and *B* values could take values between 0 and 1. Negatives values were obtained for *R* when the R primary was combined with the test stimulus.

### Normalization and transformation

Since the quantities of the test wavelengths are classically referred to an equal energy spectrum (Brainard & Stockman, 2010; Smith & Pokorny, 2003) and our test stimuli had different luminance for different wavelengths, the maximums were normalized to one, in order to refer the matching data to an EES test light (see Supplementary material for details).

In order to show our results in *SML* systems, a transformation matrix can be computed, such that:

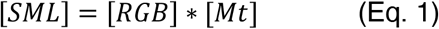

To obtain this matrix we used the averaged values of the four participants interpolated to obtain values for every 1 nm for the “Large Central” condition, which was equivalent in size to Stiles and Burch’s condition (1959). Transformation matrices are shown in Table 1 for Smith & Pokorny’s fundamentals (1975) and in Table 2 for CIE fundamentals (2006). Note that values at Tables 1 and 2 are different in terms of scale, due to the different normalization procedure for each set of fundamentals.

**Table 1.** Transformation matrix (Mt) computed to transform our data to Smith & Pokorny’s 10° fundamentals

|  | S | M | L |
| --- | --- | --- | --- |
| R | -0.5261 | 1.8261 | 13.8136 |
| G | 0.6863 | 8.9103 | 12.2892 |
| B | 77.9245 | 5.8297 | 7.0949 |

**Table 2.**
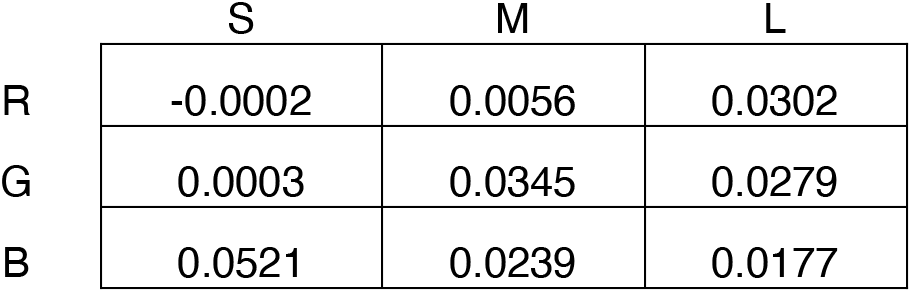
Transformation matrix (Mt) computed to transform our data to CIE 10° fundamentals

### Statistical Analysis

Repeated measures ANOVAs, Bonferroni post-hoc analyses and variance-comparison tests were used to compare color matching data among visual field conditions. Simple linear regression analysis was employed to assess data agreement with CMFs. Statistical analysis and modelling were performed in STATA 12.0 (StataCorp LP, College Station, TX).

## Results

Color matching data obtained using our primaries after normalization is shown in Figure 2 (top row). For all participants and for all conditions a similar pattern was obtained for the matches (individual data are presented in Fig. A2 of the supplementary material). As expected, for the shortest test wavelength tested (470 nm), contribution of *B* primary was larger than those of both *G* and *R* primaries to obtain a good match. For intermediate wavelengths (480 nm – 560 nm), participants needed more intensity of *G* primary and less of both *R* and *B* primaries for matching. Relatively similar contributions of R and G primary with no contribution of *B* primary were obtained for 570nm and 580 nm. For test wavelengths higher than 580 nm, R primary was predominant in the matching.

**Fig. 2.**
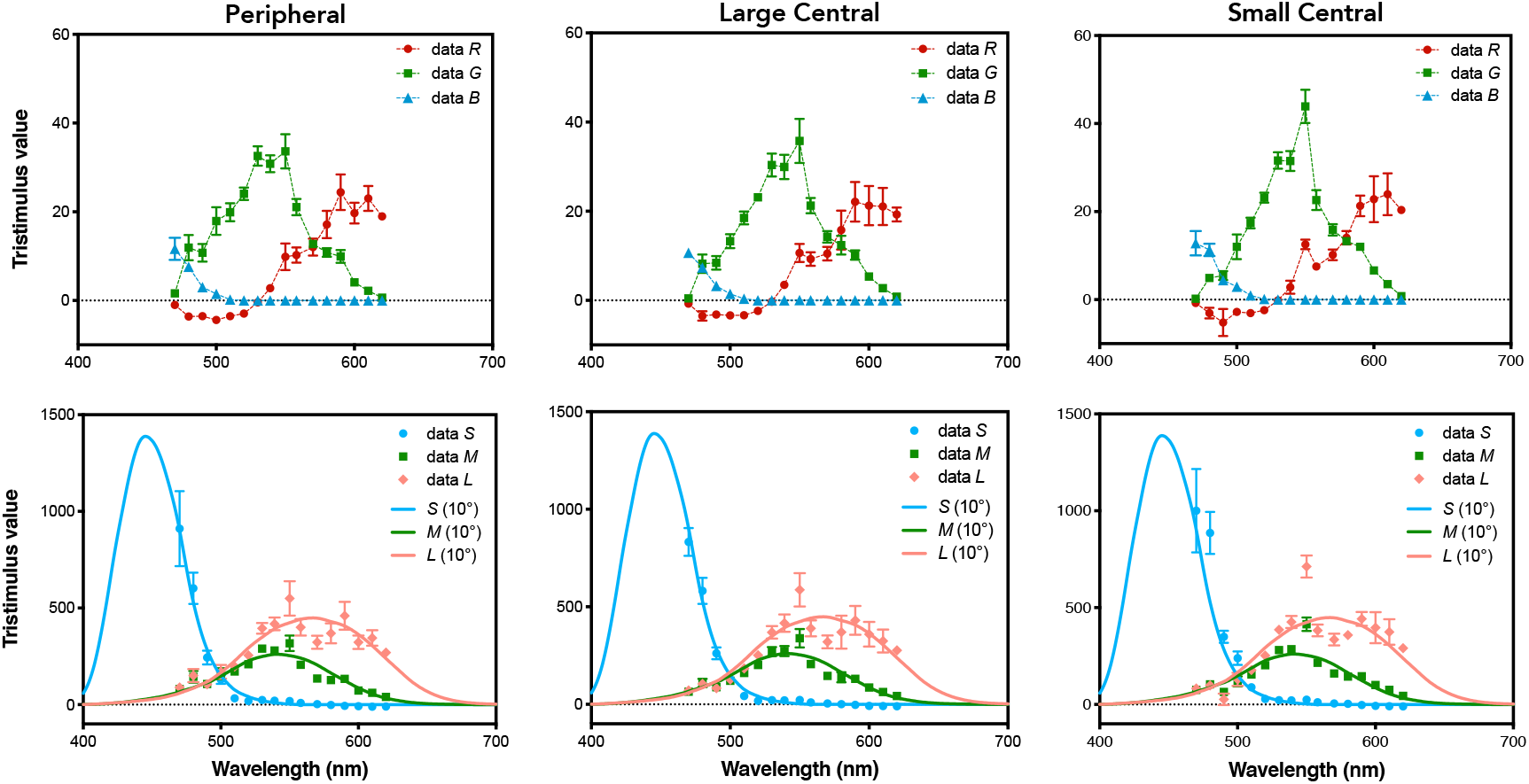
Data for three visual field conditions. Averaged color matching data of the four participants previous (top row) and after (bottom row) transformation to SML primaries. Error bars are SEM.

While *R* and *G* tristimulus values are comparable among the three visual field conditions [*R*: *F* (2, 96) = 0.05, *p* = 0.955; *G*: *F* (2, 96) = 0.77, *p* = 0.467] a significant difference was found for *B* data between the Small Central condition versus both Peripheral and Large Central conditions [*F* (2, 96) = 6.33, *p* < 0.01; Peripheral vs Small Central: *t* = 2.94, *p* < 0.05; Large Central vs Small Central: *t* = 3.21, *p* < 0.01; Large Central vs Peripheral: *t* = 0.27, *p* = 0.99]. A variability analysis showed that *R* and *G* data set have higher variability than *B* data (Supplementary Material).

Bottom panels of Figure 2 show averaged tristimulus values after transformation in *SML* system using the transformation matrix showed in Table 1, together with Smith & Pokorny’s 10° fundamentals. Linear regression analyses indicated that the S, M and L data for Large Central and Peripheral conditions were better described by the 10° cone fundamentals than the Small Central condition (Table 3). Instead, the Small Central data is better described by 2° cone fundamentals. Same conclusions applied for CIE fundamentals, although slightly better correlations were found with Smith & Pokorny’s fundamentals for L results.

**Table 3.** Adjusted-R^2^ values for comparison of our data with fundamentals from Smith & Pokorny (1975) and CIE (2006).

| Field size conditions | Smith & Pokorny's fundamentals |  |  | CIE fundamentals |  |  |
| --- | --- | --- | --- | --- | --- | --- |
|  | S | M | L | S | M | L |
| Peripheral vs 10° | 0.99 | 0.82 | 0.79 | 0.99 | 0.82 | 0.77 |
| Large Central vs 10° | 0.99 | 0.84 | 0.80 | 0.99 | 0.84 | 0.78 |
| Small Central vs 10° | 0.95 | 0.74 | 0.69 | 0.95 | 0.73 | 0.67 |
| Small Central vs 2° | 0.97 | 0.76 | 0.72 | 0.96 | 0.76 | 0.70 |

Results for the bleaching experiment are shown in Figure 3. This figure also contains the results reported above for the Peripheral condition and the Styles & Burch’s *R*, *G*, and *B* functions. There are no significant differences between Peripheral and Bleaching data [*R*: *F* (1, 36) = 0, *p* = 0.99; *G*: *F* (1, 36) = 0.33, *p* = 0.59; *B*: *F* (1, 36) = 0.1, *p* = 0.76], indicating minimal rod intrusion in the data.

**Fig. 3.**
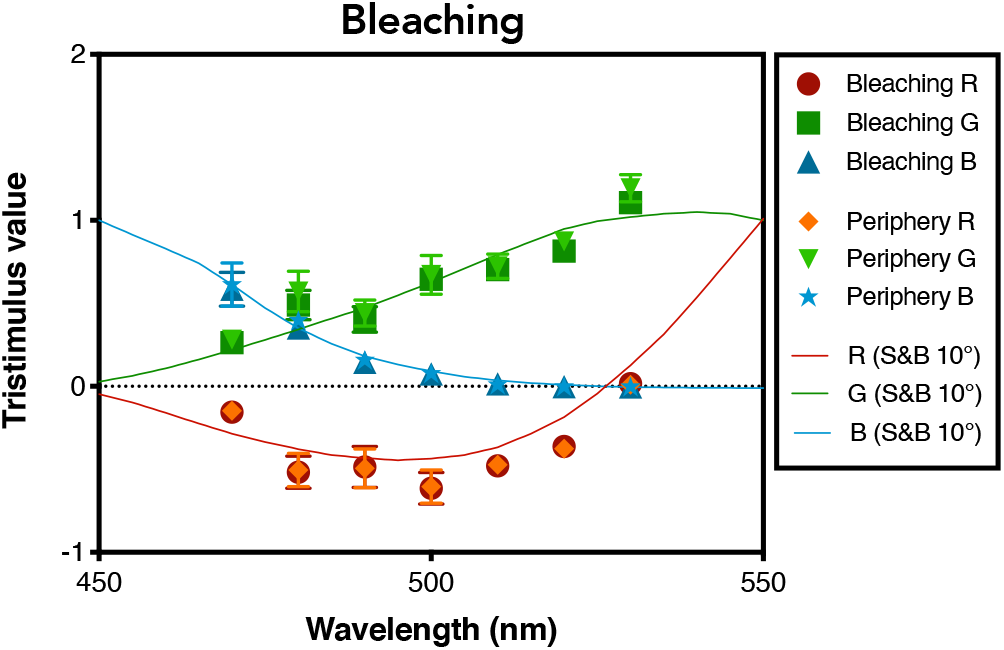
Matching data for peripheral condition after exposition to bleaching light, in the range of 470 nm – 530 nm. Bleaching data is compared with data for similar field condition of Figure 2 (left column). Error bars are SEM.

### Fundamentals modelling

A statistical analysis was performed comparing 2° and 10° Smith & Pokorny (1975) or CIE (2006) *L*, *M* and *S* CMFs. The analysis was aimed to predict 10° CMF values based on 2° CMF values (which is largely driven by cone inputs) to assess whether rod and/or melanopsin activation is necessary. To avoid over sampling, the analysis only included wavelengths every 10 nm (i.e. 31 data points). Four nested regression models were assessed with likelihood ratio test for comparison of model fits.

The first model (M_1_) assumed the contribution of rods and melanopsin to the testing condition A for predicting testing condition B (Equation 2, cone + melanopsin + rod). The second model (M_2_) assumed only the contribution of melanopsin together with results from the testing condition A (Equation 3, cone + melanopsin). The third model (M_3_) assumed only the contribution of rods together with results from the testing condition A (Equation 4, cone + rod). The fourth model (M_4_) assumed on the only contribution of with results from the testing condition A (Equation 5, cone).

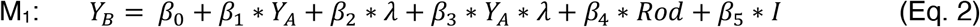

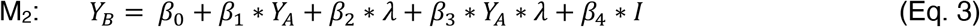

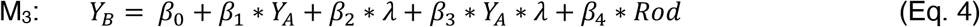

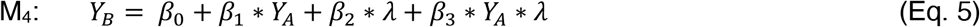

Where *Y_B_* is either *L*, *M* or *S* 10° sensitivity function, *β*_0-5_ are coefficients, *Y*_A_ is either *L*, *M* or *S* 2° sensitivity function, *λ* is wavelength in nm, rod sensitivity function *(Rod)* is based on the scotopic sensitivity function (CIE, 1951), and *I* is computed according to the melanopic spectral sensitivity function (Enezi et al., 2011), and *S*, *M, L, Rod* and *I* functions were normalized in a similar fashion as the relative Troland (Td) space (Barrionuevo & Cao, 2014; Smith & Pokorny, 1996). For CIE fundamentals, *Rod* and *I* were obtained from CIE (2018), and *S*, *M*, *L*, *Rod* and *I* functions were normalized to the maximum following CIE recommendation (CIE, 2018).

Table 4 shows the likelihood ratio test results comparing the goodness-of-fits. For *S* and *M* fundamentals M_1_ is better model than M_2_, M_3_ and M_4_. For *L* fundamental M_1_ doesn’t improve fitting compared with both M_2_ and M_3_, but it is better than M_4_.

**Table 4.**
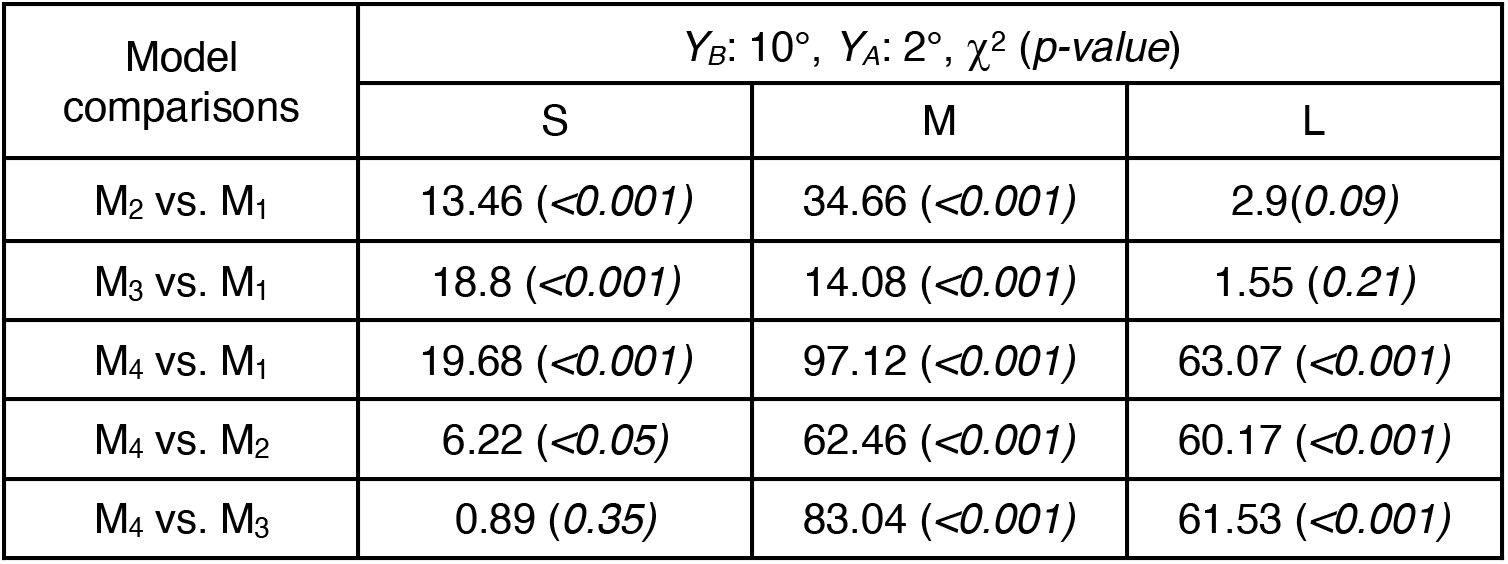
Likelihood ratio test parameters for prediction of Smith & Pokorny’s 10° fundamentals from combined effects of 2° fundamentals with Rod and Melanopsin.

| Model comparisons | $Y_B: 10^\circ, Y_A: 2^\circ, \chi^2 (p\text{-value})$ | | |
| --- | --- | --- | --- |
|  | S | M | L |
| $M_2$ vs. $M_1$ | 13.46 ($<0.001$) | 34.66 ($<0.001$) | 2.9(0.09) |
| $M_3$ vs. $M_1$ | 18.8 ($<0.001$) | 14.08 ($<0.001$) | 1.55 (0.21) |
| $M_4$ vs. $M_1$ | 19.68 ($<0.001$) | 97.12 ($<0.001$) | 63.07 ($<0.001$) |
| $M_4$ vs. $M_2$ | 6.22 ($<0.05$) | 62.46 ($<0.001$) | 60.17 ($<0.001$) |
| $M_4$ vs. $M_3$ | 0.89 (0.35) | 83.04 ($<0.001$) | 61.53 ($<0.001$) |

These analyses suggest that both rod and melanopsin might help to improve the fitting only in *S* and *M* Smith & Pokorny’s fundamentals. Interestingly, when we compared M4 and M3, there is no improvement of fitting for the *S* fundamental, suggesting no intrusion of rods.

Table 5 shows the likelihood ratio tests parameters for model comparison considering CIE fundamentals (CIE, 2006). For *S* fundamental M_1_ model is better than M_2_, M_3_ and M_4_. For *M* fundamental M_1_ is better than M_2_ and M_4_ and similar than M_3_. For *L* fundamental M_1_ is better than both M_3_ and M_4_, and similar than M_2_. This analysis suggests that for *S* fundamental both rod and melanopsin might help to improve the fitting. For *M* fundamental, melanopsin helped to improve the fitting while for *L* fundamental, rod helped to improve the fitting. Consistent with Smith and Pokorny’s fundamentals analysis, when we compared M_4_ and M_3_, there is no an improvement of fitting for *S* fundamental, suggesting no intrusion of rods.

**Table 5.** Likelihood ratio test parameters for prediction of CIE 10° fundamentals from combined effects of 2° fundamentals with Rod and Melanopsin.

| Model comparisons | $Y_B: 10^\circ, Y_A: 2^\circ, \chi^2 (p\text{-value})$ | | |
| --- | --- | --- | --- |
|  | S | M | L |
| $M_2$ vs. $M_1$ | 15.97 (<0.001) | 15.37 (<0.001) | 0.63 (0.43) |
| $M_3$ vs. $M_1$ | 20.25 (<0.001) | 3.48 (0.06) | 7.16 (<0.05) |
| $M_4$ vs. $M_1$ | 20.66 (<0.001) | 70.15 (<0.001) | 47.14 (<0.001) |
| $M_4$ vs. $M_2$ | 4.69 (<0.05) | 54.78 (<0.001) | 46.5 (<0.001) |
| $M_4$ vs. $M_3$ | 0.41 (0.52) | 66.67 (<0.001) | 39.97 (<0.001) |

### Data modelling

In the CIE (2006) recommendations, the differences between 10° cone fundamentals and 2° cone fundamentals are explained by optical density differences due to macular pigment and cone photopigments (Stockman, 2019). In order to analyze if this approach can account for the differences found in our data, we computed 10° LMS fundamentals based on our Small Central data (in CIE system) weighted by a factor that related CIE 10° fundamentals with CIE 1.7° fundamentals.

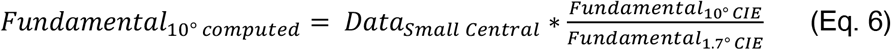

Using likelihood ratio tests and considering the models 1 – 4 (Equations 2 -5), the computed 10° Fundamentals (*Y*_A_) were compared with the Large Central data (*Y*_B_) after transformation to the CIE system and assessing the effect of rod and melanopsin intrusion. If only optical media can account for the differences between Small Central and Large Central data, then the computed 10° Fundamentals would be equal to Large Central data and all models would provide similar goodness-of-fits.

Table 6 shows the likelihood ratio tests parameters for model comparison considering CIE fundamentals (CIE, 2006). For the three datasets M_1_ model is better than M_2_, M_3_ and M_4_. However, M_2_ and M_3_ didn’t improve the fitting in comparison to M_4_. This analysis suggests that variation of intraocular media is not enough to explain differences in our results and both rod and melanopsin help to improve the fitting but not in independently.

**Table 6.** Likelihood ratio test parameters for prediction of Large Central data from combined effects of computed 10°fundamentals with Rod and Melanopsin.

| Model comparisons | $Y_B$: Large Central data, $Y_A$: 10° computed, $\chi^2$ (p-value) | | |
| --- | --- | --- | --- |
|  | S | M | L |
| $M_2$ vs. $M_1$ | 7.96 (<0.001) | 12.35 (<0.001) | 13.71 (<0.001) |
| $M_3$ vs. $M_1$ | 6.5 (<0.05) | 11.85 (<0.001) | 15.53 (<0.001) |
| $M_4$ vs. $M_1$ | 8.08 (<0.05) | 13.47 (<0.01) | 15.6 (<0.001) |
| $M_4$ vs. $M_2$ | 0.12 (0.73) | 1.12 (0.29) | 1.9 (0.19) |
| $M_4$ vs. $M_3$ | 1.58 (0.21) | 1.62 (0.2) | 0.07(0.79) |

## Discussion

We obtained color matching data for three stimulation size conditions. One involving only cones (Small Central condition) and the other two (Large Central and Peripheral conditions) with potential intrusion of rods and melanopsin. We found differences only for the *B* component when comparing results for the Large Central or the Peripheral conditions with the Small Central condition.

From a bleaching experiment, which minimized rod contribution, we did not find differences between bleaching and no-bleaching data in peripheral condition, suggesting that the differences for our data between the Small Central condition and the Peripheral are not explained by rod intrusion.

On the other hand, we conducted statistical analyses considering classical Smith & Pokorny’s (1975) and CIE (2006) *L*, *M* and *S* fundamentals. From these analyses we found that the difference between 10° and 2° conditions is explained by a combination of rods and melanopsin for *S* and *M* Smith & Pokorny’s fundamentals and *S* CIE fundamental. Also, the statistical model on CIE fundamentals for explaining differences needed to incorporate only melanopsin for the *M* fundamental and only rods for the *L* fundamental. Furthermore, the inclusion of melanopsin to the only cone model improved the fitting of the *S* fundamental, but the inclusion of rods did not. From this approach, it is clear that melanopsin may play a role in the large field sensitivity functions, especially for the *S* fundamental.

An alternative hypothesis is that the differences between large fields and small fields CMFs were accounted only for eccentrical variation of macular filtering and subtle peak density change of visual pigments (CIE, 2006; Stockman, 2019). Incorporation of the effect of these filters to our data didn’t explain completely the differences between Small Central and Large Central conditions. Instead, incorporation of the filters effect in addition with rods and melanopsin produced a better fit.

The need of CIE 10° CMFs recommendations was consequence of recognizing that color matches are field size dependent. All the analysis and experiments that we conducted suggested a role of melanopsin in large field CMFs. However, we realized that this study is limited, and to better quantify the participation of melanopsin in CMFs, further experiments with more participants need to be carried out.

The agreement between our results for the Large Central and Peripheral conditions indicates that the observers ignored the perception of central area in Large Central condition to produce the judgement, since the ring-shape of the Peripheral configuration avoided light stimulation of the Maxwell spot. This conclusion is in agreement with statements on previous studies (Stiles & Burch, 1959; Stockman, 2019). It gives a hint that for large field stimulus, color judgements could be mostly driven by extrafoveal perception, which stressed the importance to understand the components in the peripheral color vision.

Classical color matching procedure consists in presenting the test and matching stimuli simultaneously forming a bipartite field, therefore, the perceptual matching is achieved from a spatial comparison. A potential drawback of this configuration is that the stimuli are imaged at different parts of the retina, which constitutes an asymmetric comparison (Wyszecki & Stiles, 2000). We presented our stimuli in a sequential manner with slow alternation. We decided to use this configuration to image both stimuli at the same retinal area for activation of the same photoreceptors. Our data are very similar to those obtained with the bipartite field, which can be seen as a mutual validation of both techniques.

Due to the sluggish nature of melanopsin, it is possible that in regular conditions with natural saccadic movements, the effect of melanopsin in peripheral color perception might be insignificant and trichromacy holds. However, from our results there might be a role of melanopsin activation in CMFs. Therefore, the current color specification recommended by CIE might be compromised in extrafoveal conditions and for short wavelengths.

## References

Aranda, M. L., & Schmidt, T. M. (2020). Diversity of intrinsically photosensitive retinal ganglion cells: Circuits and functions. Cellular and Molecular Life Sciences. https://doi.org/10.1007/s00018-020-03641-5

Barrionuevo, P. A., & Cao, D. (2014). Contributions of rhodopsin, cone opsins, and melanopsin to postreceptoral pathways inferred from natural image statistics. Journal of the Optical Society of America A, 31(4), A131–A139. https://doi.org/10.1364/JOSAA.31.00A131

Barrionuevo, P. A., & Cao, D. (2019). Does melanopsin help to explain color constancy in natural environments? In Proceedings of the International Color Association (AIC) Conference 2019 (pp. 598–605). International Colour Association Incorporated.

Berson, D. M., Dunn, F. A., & Takao, M. (2002). Phototransduction by Retinal Ganglion Cells That Set the Circadian Clock. Science, 295(5557), 1070–1073. https://doi.org/10.1126/science.1067262

Brainard, D. H., & Stockman, A. (2010). Colorimetry. In Handbook of Optics: Vol. III Vision and Vision Optics. McGraw Hill.

Brown, T. M., Tsujimura, S., Allen, A. E., Wynne, J., Bedford, R., Vickery, G., Vugler, A., & Lucas, R. J. (2012). Melanopsin-Based Brightness Discrimination in Mice and Humans. Current Biology, 22(12), 1134–1141. https://doi.org/10.1016/j.cub.2012.04.039

Cao, D., Chang, A., & Gai, S. (2018). Evidence for an impact of melanopsin activation on unique white perception. JOSA A, 35(4), B287–B291. https://doi.org/10.1364/JOSAA.35.00B287

Cao, D., Nicandro, N., & Barrionuevo, P. A. (2015). A five-primary photostimulator suitable for studying intrinsically photosensitive retinal ganglion cell functions in humans. Journal of Vision, 15(1), 27. https://doi.org/10.1167/15.1.27

CIE. (1932). Commission Internationale de l’Eclariage Proceedings, 1931. Cambridge University Press.

CIE. (1951). Commission Internationale de l’Eclariage Proceedings, 1951. Bureau Central de la CIE.

CIE. (1964). Proceedings, Vienna Session, 1963: Vol. B. Bureau Central de la CIE.

CIE. (2006). Fundamental chromaticity diagram with physiological axes—Part 1 (CIE 170-1:2006). http://cie.co.at/publications/fundamental-chromaticity-diagram-physiological-axes-part-1

CIE. (2018). *CIE SYSTEM FOR METROLOGY OF OPTICAL RADIATION FOR IPRGC-INFLUENCED RESPONSES TO LIGHT* (International Standard S 026/E:2018). International Commission on Illumination. https://doi.org/10.25039/S026.2018

Dacey, D. M., Liao, H.-W., Peterson, B. B., Robinson, F. R., Smith, V. C., Pokorny, J., Yau, K.-W., & Gamlin, P. D. (2005). Melanopsin-expressing ganglion cells in primate retina signal colour and irradiance and project to the LGN. Nature, 433(7027), 749–754. https://doi.org/10.1038/nature03387

Enezi, J. al, Revell, V., Brown, T., Wynne, J., Schlangen, L., & Lucas, R. (2011). A “Melanopic” Spectral Efficiency Function Predicts the Sensitivity of Melanopsin Photoreceptors to Polychromatic Lights. Journal of Biological Rhythms, 26(4), 314–323. https://doi.org/10.1177/0748730411409719

Foster, R. G., Hughes, S., & Peirson, S. N. (2020). Circadian Photoentrainment in Mice and Humans. Biology, 9(7), 180. https://doi.org/10.3390/biology9070180

Geisler, W. S., & Banks, M. S. (2010). Visual performance. In M. Bass (Ed.), Handbook of optics: Vol. III Vision and Vision Optics (3rd ed., p. 2.1–2.51). The McGraw-Hill Companies Inc.

Hattar, S., Liao, H.-W., Takao, M., Berson, D. M., & Yau, K.-W. (2002). Melanopsin-Containing Retinal Ganglion Cells: Architecture, Projections, and Intrinsic Photosensitivity. Science, 295(5557), 1065–1070. https://doi.org/10.1126/science.1069609

Hattar, S., Lucas, R. J., Mrosovsky, N., Thompson, S., Douglas, R. H., Hankins, M. W., Lem, J., Biel, M., Hofmann, F., Foster, R. G., & Yau, K.-W. (2003). Melanopsin and rod–cone photoreceptive systems account for all major accessory visual functions in mice. Nature, 424(6944), 75–81. https://doi.org/10.1038/nature01761

Horiguchi, H., Winawer, J., Dougherty, R. F., & Wandell, B. A. (2013). Human trichromacy revisited. Proceedings of the National Academy of Sciences, 110(3), E260–E269. https://doi.org/10.1073/pnas.1214240110

Lucas, R. J., Allen, A. E., Milosavljevic, N., Storchi, R., & Woelders, T. (2020). Can We See with Melanopsin? Annual Review of Vision Science, 6(1), null. https://doi.org/10.1146/annurev-vision-030320-041239

Moreland, J. D., & Cruz, A. (1959). Colour Perception with the Peripheral Retina. Optica Acta: International Journal of Optics, 6(2), 117–151. https://doi.org/10.1080/713826278

Nakano, Y. (1996). Part III: Color vision mathematics: A tutorial. Human Color Vision, 544–562.

Palmer, D. A. (1981). Nonadditivity in color matches with four instrumental stimuli. JOSA, 71(8), 966–969. https://doi.org/10.1364/JOSA.71.000966

Provencio, I., Rodriguez, I. R., Jiang, G., Hayes, W. P., Moreira, E. F., & Rollag, M. D. (2000). A Novel Human Opsin in the Inner Retina. The Journal of Neuroscience, 20(2), 600–605.

Smith, V. C., & Pokorny, J. (1975). Spectral sensitivity of the foveal cone photopigments between 400 and 500 nm. Vision Research, 15(2), 161–171. https://doi.org/10.1016/0042-6989(75)90203-5

Smith, V. C., & Pokorny, J. (1996). The design and use of a cone chromaticity space: A tutorial. Color Research & Application, 21(5), 375–383. https://doi.org/10.1002/(SICI)1520-6378(199610)21:5<375::AID-COL6>3.0.CO;2-V

Smith, V. C., & Pokorny, J. (2003). Color matching and color discrimination. The Science of Color, 2003, 103–148.

Spitschan, M. (2019). Photoreceptor inputs to pupil control. Journal of Vision, 19(9), 5–5. https://doi.org/10.1167/19.9.5

Stiles, W. s., & Burch, J. m. (1959). N.P.L. Colour-matching Investigation: Final Report (1958). Optica Acta: International Journal of Optics, 6(1), 1–26. https://doi.org/10.1080/713826267

Stockman, A. (2019). Cone fundamentals and CIE standards. Current Opinion in Behavioral Sciences, 30, 87–93. https://doi.org/10.1016/j.cobeha.2019.06.005

Trezona, P. W. (1970). Rod participation in the ‘blue’ mechanism and its effect on colour matching. Vision Research, 10(4), 317–332. https://doi.org/10.1016/0042-6989(70)90103-3

Wyszecki, G., & Stiles, W. S. (2000). Color Science: Concepts and Methods, Quantitative Data and Formulae. Wiley.

Yamakawa, M., Tsujimura, S., & Okajima, K. (2019). A quantitative analysis of the contribution of melanopsin to brightness perception. Scientific Reports, 9(1), 7568. https://doi.org/10.1038/s41598-019-44035-3

Zele, A. J., Adhikari, P., Feigl, B., & Cao, D. (2018). Cone and melanopsin contributions to human brightness estimation. JOSA A, 35(4), B19–B25. https://doi.org/10.1364/JOSAA.35.000B19

Zele, A. J., Feigl, B., Adhikari, P., Maynard, M. L., & Cao, D. (2018). Melanopsin photoreception contributes to human visual detection, temporal and colour processing. Scientific Reports, 8(1), 3842. https://doi.org/10.1038/s41598-018-22197-w

