## Supplementary material for "Is Melanopsin Activation Affecting Large Field Color Matching Functions?"

Explanation of the normalization to EES

Since the quantities of the test wavelengths are customarily referred to an equal energy spectrum (Smith & Pokorny, 2003; Brainard & Stockman, 2010) and our test lights have different energies ($E_i$), which is evident from the filter transmittance profile of Figure 1, we weighted the $R_i$, $G_i$ and $B_i$ results considering the maximums for each test light (*i*-th). This weight ($k_i$) shouldn't affect the equality, since:

$$\frac{1}{k_i}(R_i + G_i + B_i) = \frac{1}{k_i}E_i$$

This normalization was conducted to follow the literature. An example of the effect of the normalization on the data is shown in Fig. A1.

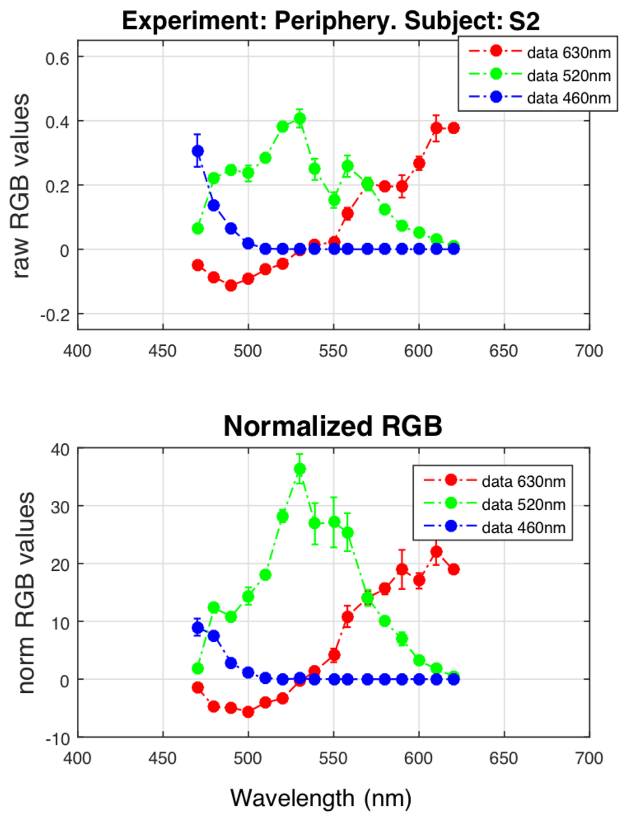


Fig. A1. RGB data from subject S2 before (upper panel) and after (lower panel) normalization for the Peripheral Condition.

Individual data of figure 2.

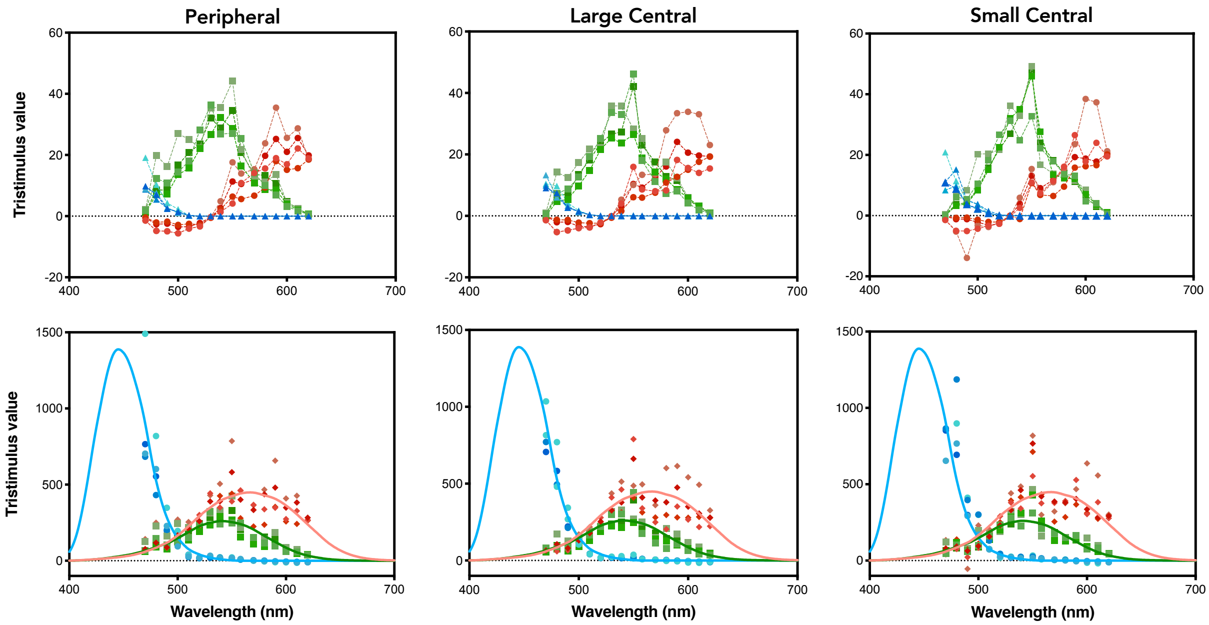


Fig. A2. Data for three visual field conditions. Individual color matching data of the four participants before (top row) and after (bottom row) transformation to SML primaries.

Variability Comparison of *R*, *G* and *B* data

In the three visual field conditions, variability between participants was similar for *R* and *G* values (Peripheral: $f = 0.9495$, $p = 0.8376$; Large Central: $f = 0.9122$, $p = 0.7163$; Small Central: $f = 0.7667$, $p = 0.2944$), and both *R* and *G* data sets have higher variability than *B* values [data *R* vs data *B* (Peripheral: $f = 9.8084$, $p < 0.05$; Large Central: $f = 11.7818$, $p < 0.05$; Small Central: $f = 6.6712$, $p < 0.05$); data *G* vs data *B* (Peripheral: $f = 10.3303$, $p < 0.05$; Large Central: $f = 12.9164$, $p < 0.05$; Small Central: $f = 8.7012$, $p =< 0.05$)].
